# Quantifying the hydrological niche of swamp vegetation communities using indicator species

**DOI:** 10.1101/2025.05.18.653618

**Authors:** David C. Deane, Tanya Mason, Joe Cairns, Martin Krogh, David Keith

## Abstract

Indicator species are typically used to infer the presence of other biota or specific environmental conditions. Here we use indicator species to quantify the hydrological niche of their corresponding vegetation communities within Coastal Upland Swamps of the Sydney Basin Bioregion, Australia. Swamp vegetation organizes naturally into five recognised communities, thought to occupy distinct positions along a hydrological gradient. However, longwall mining reduces hydro-period, potentially impacting the observed vegetation. We modelled the hydrological niche for each community using the relative frequency of 20 vascular plant indicator species (four for each community) and time series soil moisture data from 11 unmined sites across four swamps. We then used this model to predict indicator species frequencies in 3 mine-impacted sites. Indicator species were modelled as a function of the average number of days-per-year that swamps remained saturated at the base of the root zone (30 cm below surface). Hydrological niches were well differentiated for four communities with the estimated optimal mean annual days-per-year saturated ranging from Restioid heath at 16 [<16, 29] (mean ± [95% uncertainty]) to Ti-tree thicket at 352 [322, >353]. Banksia thicket 96 [65, 154] and Cyperoid heath 257 [204, 297] communities were intermediate. However, Sedgeland indicator species showed limited variation with changing saturation, and their hydrological niche remains unclear. The model under-predicted the frequency of Cyperoid heath and over-predicted Banksia thicket indicator species in mine-impacted sites, suggesting vegetation is not yet in equilibrium with hydrology. Results suggest indicator species can provide a reliable basis for determining the hydrological niche of wetland plant communities, which can in turn predict community-level impacts of hydrological change.

## Introduction

Freshwater biodiversity faces elevated extinction risk from multiple stressors (Sayer et al. 2025), which is expected to increase further in future (Dudgeon 2019). Relative to their area, wetlands contain disproportionately high global freshwater biodiversity (Dudgeon et al. 2006, Vörösmarty et al. 2010), yet threats to many wetland-dependent taxa remain poorly understood (Sayer et al. 2025). Accordingly, improved understanding of the cause-and-effect relationships between wetland biodiversity and ecological stressors is a research priority (Birk et al. 2020, Maasri et al. 2022). Among these causal relationships, predicting relative risk of impacts from altered hydrological regimes—and thus the extent to which wetlands could provide future refugia for biodiversity—are among the most pressing questions (Taylor et al. 2021). However, a major challenge to quantifying hydrological cause-effect relationships for wetland biodiversity is the degree of natural variation within and among freshwater ecosystems (Norris et al. 2012, Campbell et al. 2022).

To simplify the task of quantifying ecological cause-effect relationships in wetlands, some degree of generalization is often useful. Because hydrology exerts a dominant influence on wetland plant organisation, one can adopt a bottom-up generalization by grouping species according to their hydrological niche. Examples include the Australian water plant functional groups (Casanova and Brock 2000, Casanova 2011), which have proven useful in predicting flooding response and defining group-level hydrological niches (Campbell et al. 2014, Johns et al. 2015, Deane et al. 2017, Mason et al. 2022, Deane et al. 2025). Similarly, Ellenberg Indicator F (moisture) values (Dengler et al. 2023, Tichý et al. 2023) have been used in a comparable manner in Europe (Dwyer et al. 2021, van Willegen et al. 2025).

Alternatively, one can adopt a top-down approach, where distinct wetland plant communities are empirically defined and the associated hydrological conditions supporting that community determined (e.g., Todd et al. 2010, Gann and Richards 2015, Huang et al. 2022). Under this community-level approach, a further step toward generalization is to identify a small number of surrogate (e.g., indicator or umbrella) species, whose relative abundance is used to infer that of the broader community, and by extension, the underlying hydrological conditions upon which they depend (Siddig et al. 2016, Tälle et al. 2023). By studying the response of the smaller number of surrogate species, insights into the controls of the distribution of the broader community they represent can be obtained for a relatively low investment in time and effort, while simplifying monitoring and reporting needs (Siddig et al. 2016).

For an indicator species approach to be effective in elucidating causal relationships, there should be established ecological relationships between the surrogate (ideally multiple surrogate) species and the broader association these represent (Lindenmayer and Likens 2011, Tälle et al. 2023). In this study, we explored the utility of wetland plant indicator species to characterise different vegetation communities in an endangered upland swamp ecosystem. Our aim was to quantify the hydrological niche of each community based on the relative frequency of their indicator species and compare those inferred responses with what is known about the hydrological niche space associated with the communities. If estimated niche positions align with established wetland hydro-synecology, this would provide a line of evidence that indicator species alone can be used to quantify the hydrological niche for their community. This would not only simplify determination of community environmental water needs but also support the use of estimated hydrological niches in predicting future hydrology-driven shifts in the corresponding vegetation community.

## Methods

### Study system

Coastal Upland Swamps of the Sydney Basin Bioregion (Fig. 1), hereafter ‘Upland Swamps’, develop within poorly drained headwater valley floors or as seepage fronts on impermeable sandstone benches (Keith and Myerscough 1993). Most upland swamps occur at elevations between 200 m and 450 m (NSW Scientific Committee 2012). Swamp vegetation tends to be a treeless, fine-scale mosaic of woody shrubs or herbaceous vegetation, thought to emerge from an interplay between hydrological gradients and wildfire frequency (Keith and Myerscough 1993, Keith 1994, Keith and Tozer 2012, Mason et al. 2017). Specifically, accumulation of sediments combined with rainfall and terrestrial inflows (Young 1986) create anoxic soil conditions. Resultant vegetation communities primarily stratify along a hydrological gradient (Mason et al. 2017) such that Ti-tree thicket (dense shrub stratum and tall, dense fern and sedge stratum) and Cyperoid heath (dense large cyperaceous sedge stratum) communities occupy the wettest end of the gradient. Banksia thicket (bt), Restioid heath (rh) and Sedgeland (sl) occupy similar and drier areas within the hydrological niche space, with fire frequency and woody biomass at least partly determining their realization at a given location (Keith and Myerscough 1993). In this study, we first fit a model of all five communities but consistent with prevailing understanding, we then repeated our analysis pooling together these three drier communities (see Modelling).

**Figure 1:**
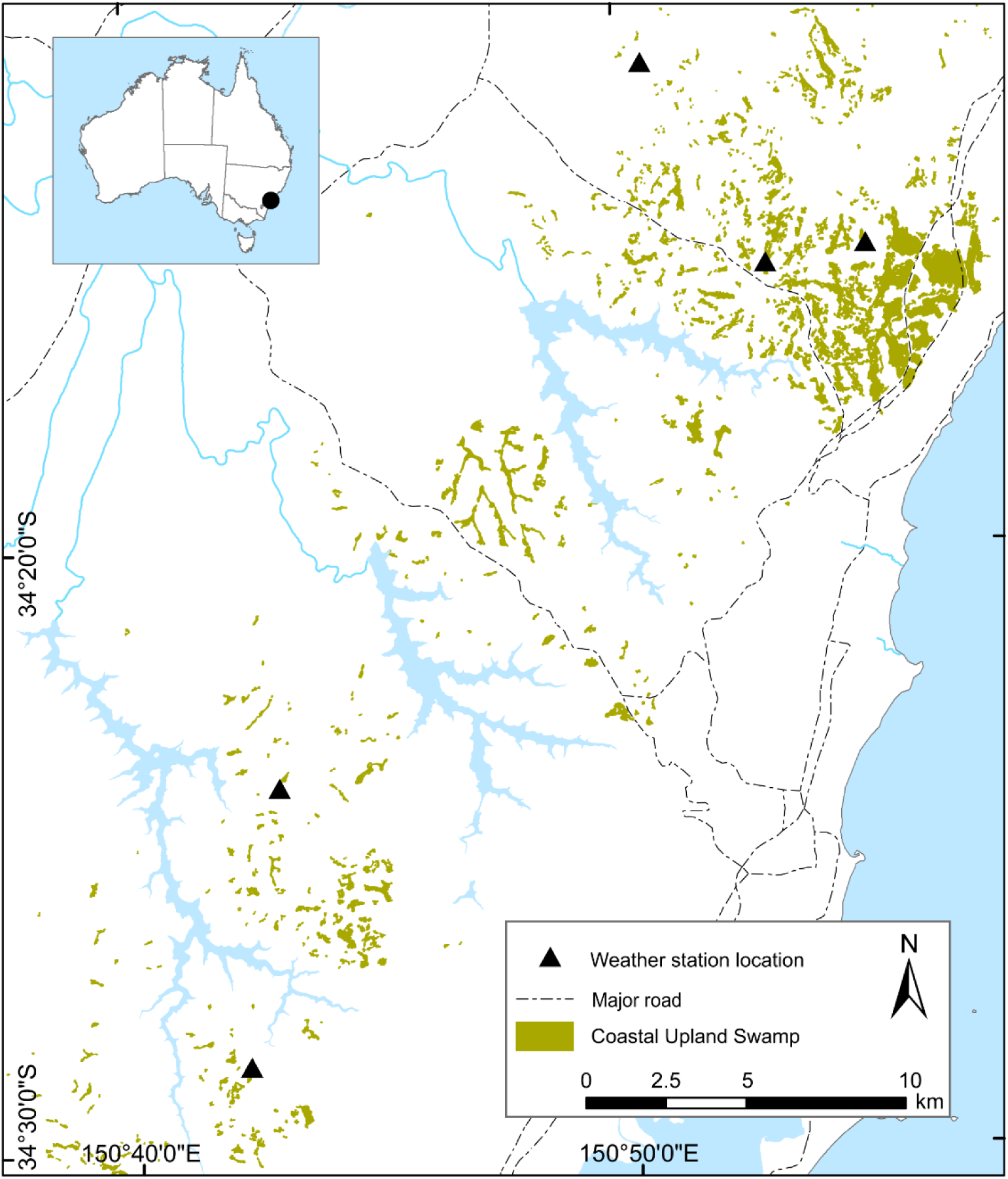
Location of weather station hydrological monitoring sites (including soil moisture probes) within the distribution of Upland Swamps. Study location within Australia is shown inset. Swamp layer obtained from Fryirs and Hose (2016).

### Indicator species for hydrological communities

Floristic classifications of five major Upland Swamp vegetation communities have been identified (Table 1). Indicator species for each community were identified using a two-way table of floristic composition (Keith and Myerscough 1993; Appendix 1) and supplemented with expert opinion to maximise detection likelihood. Here, we first fit a model of all five communities but consistent with prevailing understanding, we then repeated our analysis pooling together these three drier communities (see Modelling).

**Table 1.**
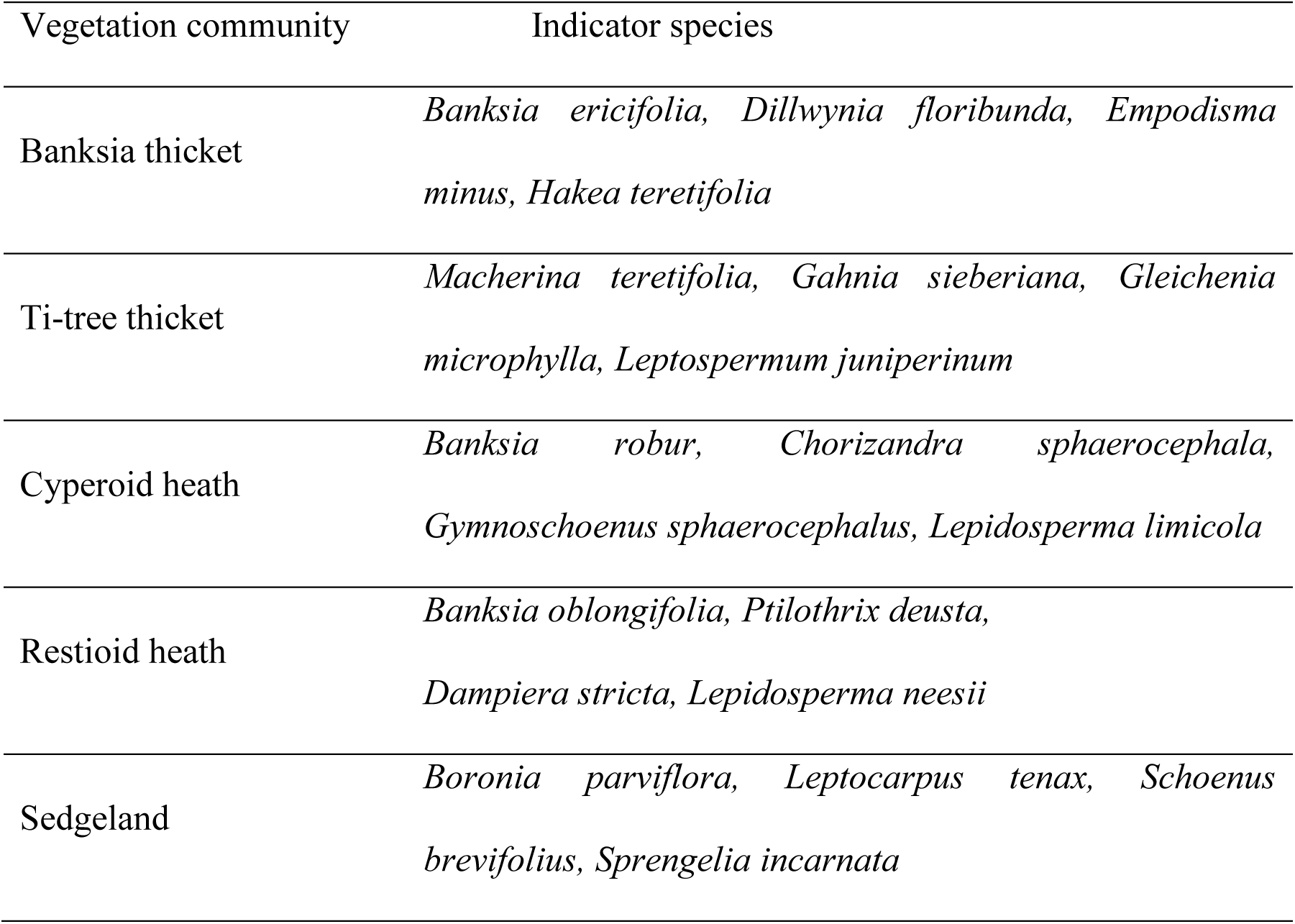
Community indicator species from Mason et al. (2021) and Keith and Myerscough (1993)

### Sources of data

We selected data from Upland Swamps included in the monitoring network described in Mason et al. (2021) that were unaffected by mining at the time of vegetation survey. Upland Swamps tend not to hold ponded surface water but are instead characterised by saturation of wetland organic peaty substrates. We therefore used data from dielectric soil moisture probes (Series DEnviroPro Dielectrics Pty Ltd., South Australia) measuring percentage saturation at four depths below ground (10, 20, 30 and 40 cm), three of which were installed in each swamp in the network. Soil moisture probes were calibrated to field soil conditions as described in Mason et al. (2021) and only sites where soil moisture probes had more than three years of data were considered here. Both soil moisture and vegetation data required pre-processing and harmonization before modelling, which is described below.

### Soil moisture data pre-processing

From a total possible number of 20 sites in the original network, we selected sites with at least 3 years of data limited to sites where upper values of soil moisture reached an asymptote, indicative of 100% saturation. This yielded 11 sites from four wetlands that had not been directly impacted by longwall mining at the time of vegetation sampling and three additional sites within one wetland where longwall mining had impacted soil moisture dynamics (Mason et al. 2021). We used only unimpacted sites to model indicator species frequency to obtain the best estimate of how unimpacted hydrological conditions result in the development of different swamp vegetation communities, while avoiding any confounded hydrological response due to time lags between the onset of hydrological change and vegetation community responses (Mason et al. 2021). We did, however, use hydrological data from the three impacted sites to predict the probability of observing the five communities using the modelled response in the unimpacted sites. By comparing the observed indicator species frequencies with the predictions from the unimpacted sites we obtained some insights into hydrology-driven changes in vegetation.

We used the 30 cm depth probe at each site, selecting this depth based on (i) its use in agricultural studies of waterlogging impacts on crop production (e.g., SEW30, see review in Liu et al. 2020), (ii) due to its use in wetland delineation (U.S. Army Corp of Engineers 1987), and (iii) based on prior work estimating the threshold of transition between wetland and terrestrial plants being within this range of sub-surface saturation (Deane et al. 2025).

Time-series of soil moisture content at 30 cm indicated clear periods of saturation of varying duration in all included sites (Appendix A, Supporting Information), but drift in the record precluded direct calculation of the proportion of time the base of the root zone was saturated (i.e., a probe reading of 100%) from the raw data. To compensate for this, soil moisture data were first de-trended and re-scaled to correct soil moisture records across the full period of record to a constant maximum reading of 100% where saturation was indicated using a moving window approach (Fig. S1.1, S1.2; Appendix A, Supporting Information). As our predictor for indicator species frequency, we calculated the mean proportion of time that the (de-trended and re-scaled) data suggested the soil profile was saturated, using 99% saturation as our threshold. Any missing days were removed from the record and the total years used in the denominator were calculated as the total number of days with observations divided by 365.

### Vegetation data pre-processing

Vegetation data corresponding to the location of soil moisture probes were collected using one of two different vegetation survey types. Each recorded species frequency (presence-absence) in 16 x 0.25 m^2^ quadrats (i.e., 0.5 x 0.5 m) but with differences in their sampling geometry and level of floristic detail. Rapid assessment data were from a 2 x 2 m quadrat divided into 16 x 0.25 m^2^ grid cells centred on the soil moisture probe location. Rapid Assessment recorded *only* the presence of the 20 indicator species (Table 1) within the sampling grid. In contrast, belt transect surveys recorded the presence of *all* vascular plant species within 0.25 m^2^ contiguous cells positioned along a 15 m linear transect starting at the probe location. To ensure a constant total surveyed area and number of cells in all surveys, we used only the first 16 cells for belt transects, which were closest to the probe location. To harmonise vegetation data, we first removed all non-indicator species recorded from the belt transect data. For both survey types, we then summed the frequency of all indicator species at each site and calculated their relative frequency as the ratio of each species frequency over the total sum of recorded frequencies for all indicator species. Thus, our response variable was the total relative frequency of indicator species for each of the five communities at each site. Analysis of model residuals indicated no influence of vegetation survey method on results (see Appendix 2).

### Modelling

To understand how soil saturation at the base of the root zone affected the relative frequency of indicator species, we used a Bayesian regression approach. To account for the spatial structure in the data, we used a multilevel (or mixed) model, with grouping terms (random intercepts) for each site nested within swamps. Models were estimated with Hamiltonian Monte Carlo (HMC) sampling using Stan (Carpenter et al. 2017) and implemented using R package brms (Burkner 2017) within the R programming environment, version 4.4.2 (R Core Team 2024). We used weakly regularizing priors and 4 independent MCMC chains, each with 2000 iterations (1000 warm-up) to estimate the model. Model convergence was verified using the potential scale reduction factor (R-hat), with all values ≤ 1.01, the bulk effective sample size (all values > 600), and by inspecting the HMC chains to ensure mixing. Residuals were then tested using R package DHARMa (Hartig 2024) (see Appendix S2 for model diagnostics). Tests on the influence of sampling method (i.e., survey type) also used model residuals and showed no systematic differences in distribution (chi-squared = 0.05, *df* = 1, *P* = 0.83; Appendix 2). Model performance was evaluated using the Bayesian R^2^ (Gelman et al. 2019).

Our response data were proportions on the half-closed interval [0, 1) and we therefore used a zero-inflated beta distribution. To account for random variation among sites and communities, we modelled the zero-inflation parameter (*zi*) modelling the mean response as an interaction between saturation and community. We modelled mean community indicator species relative frequencies (IS_frq) at each site as a function of the mean annual duration of saturation at 30 cm (Sat30), along with a quadratic term (Sat30q) to account for non-linearity in response and an indicator species community factor variable with five levels (com_IS). We also allowed indicators for each community to vary via an interaction between the linear (Sat30) and with the quadratic (Sat30q) saturation term to allow for differences in the shape and location of the response between communities. The model structure was as follows:

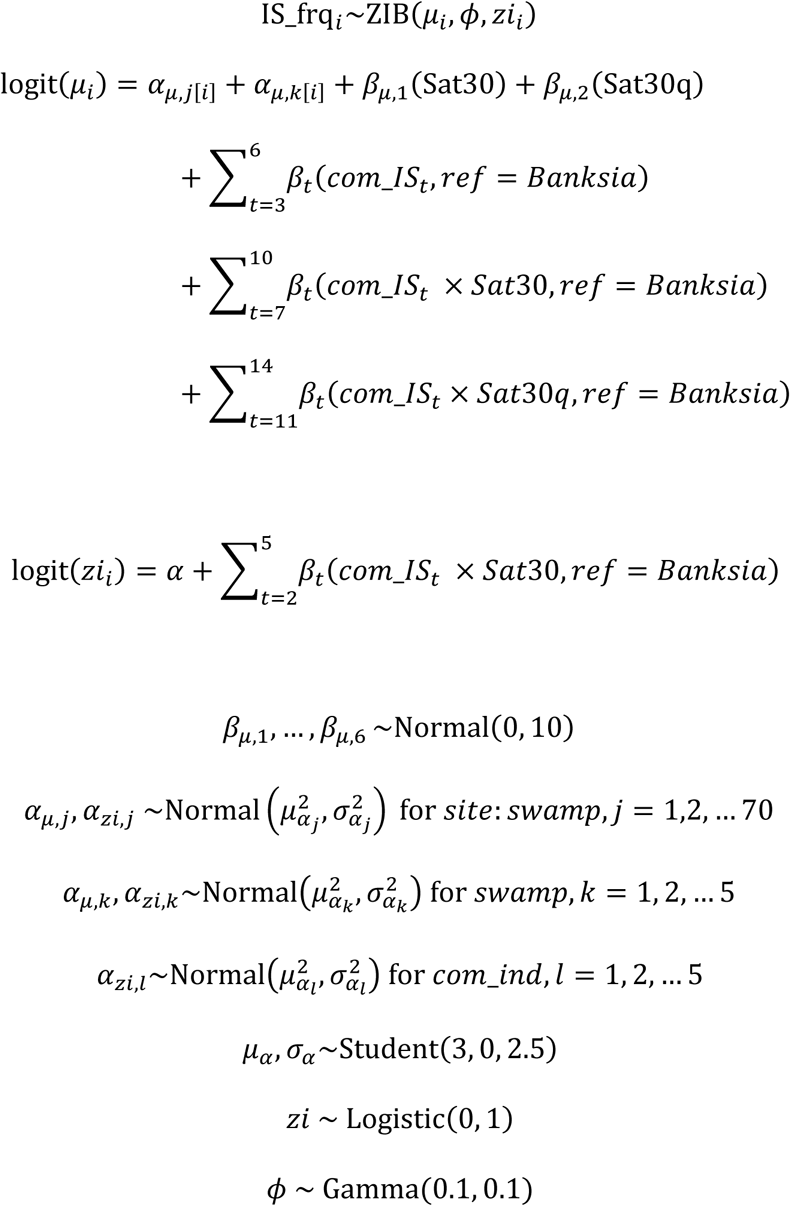

Because of the hypothesized overlap in the hydrological niche of the three communities adapted to drier conditions, we re-fitted the above model having first pooled the relative frequencies for Banksia thicket, Restioid heath and Sedgeland. In this case, the model was identical to the above, simply using three communities: the pooled *Drier composite* community, Cyperoid heath and Ti-tree thicket. We present the results for all five communities modelled separately in the main text, and those for the three communities (pooling the three dry communities) in the Supporting information.

### Post processing and prediction

We inferred the hydrological niche for each community based on the modelled variations in the corresponding community indicator species relative frequencies. We first drew 1000 samples from the posterior prediction of the final model, excluding uncertainty in the model group- and population-level variances (i.e., including uncertainty in fixed, but not random effects). The point on the *x*-axis (corresponding to mean annual proportion of time saturated) where the mean value of the posterior predictive distribution reached its maximum was taken as our estimate of the optimal hydrological niche position. To estimate variability in the estimated optima (i.e., community hydrological niche width), we calculated an uncertainty interval as the point on the *x*-axis where the mean posterior prediction reached 95% of the maximum probability. Note for the two extreme communities (i.e., the wettest and driest), this approach could not be used as predictions fell outside of the range of observed values. Given the empirical distribution of saturation values, this likely corresponds to 0 and 100% duration, but we adopt a conservative approach of referring to the corresponding uncertainty simply as being greater or less than the observed optima. For a more intuitive interpretation, we convert all written estimates of the proportion of time saturated for hydrological niches to mean days-per-year by multiplying the estimated proportions by 365.

To explore the impact of longwall mining, we used the model to predict the expected values for each indicator species group using the observed soil saturation values for the three impacted sites. We compared the observed mean (±1 SD) estimate across the three sites with the 95% credible intervals from the posterior predictive distribution (PPD). We interpreted the results by comparing the posterior predictive distribution in indicator species frequencies and the inferred mean, testing the hypothesis that the observed value – PPD was equal to zero. Thus, probabilities near 0.5 support the hypothesis that the predicted and observed were consistent, while values near 0 or 1 suggest the model under- or over-estimated observed values respectively.

## Results

Mean annual root zone saturation explained almost half of the variation in indicator species frequencies across the five communities (Bayesian R^2^ = 0.46). Response curves for the indicator species groups showed either a concave-down unimodal or broad linear response curve (Fig. 2), although the limited number of data points suggests the shape of the response should be interpreted with caution. Generally, hydrological niche responses were as hypothesized in terms of relative saturation preferred by the different communities, with Ti tree thicket preferring the wettest conditions (mean ±[95% uncertainty] = 353 [330, ≥353] days-per-year) and Cyperoid heath also preferring extended, but shorter average saturated duration (257 [204,297] d/y). Restioid heath (16 [<16, 29] d/y) and Banksia thicket (96 [65, 154] d/y) clearly preferred drier conditions. Sedgeland indicator species frequencies differed somewhat from expectation, apparently being similarly adapted to Cyperoid heath (Fig. 2) and reaching an optimal value at maximum saturation of 253 [183, ≥353] days.

**Figure 2.**
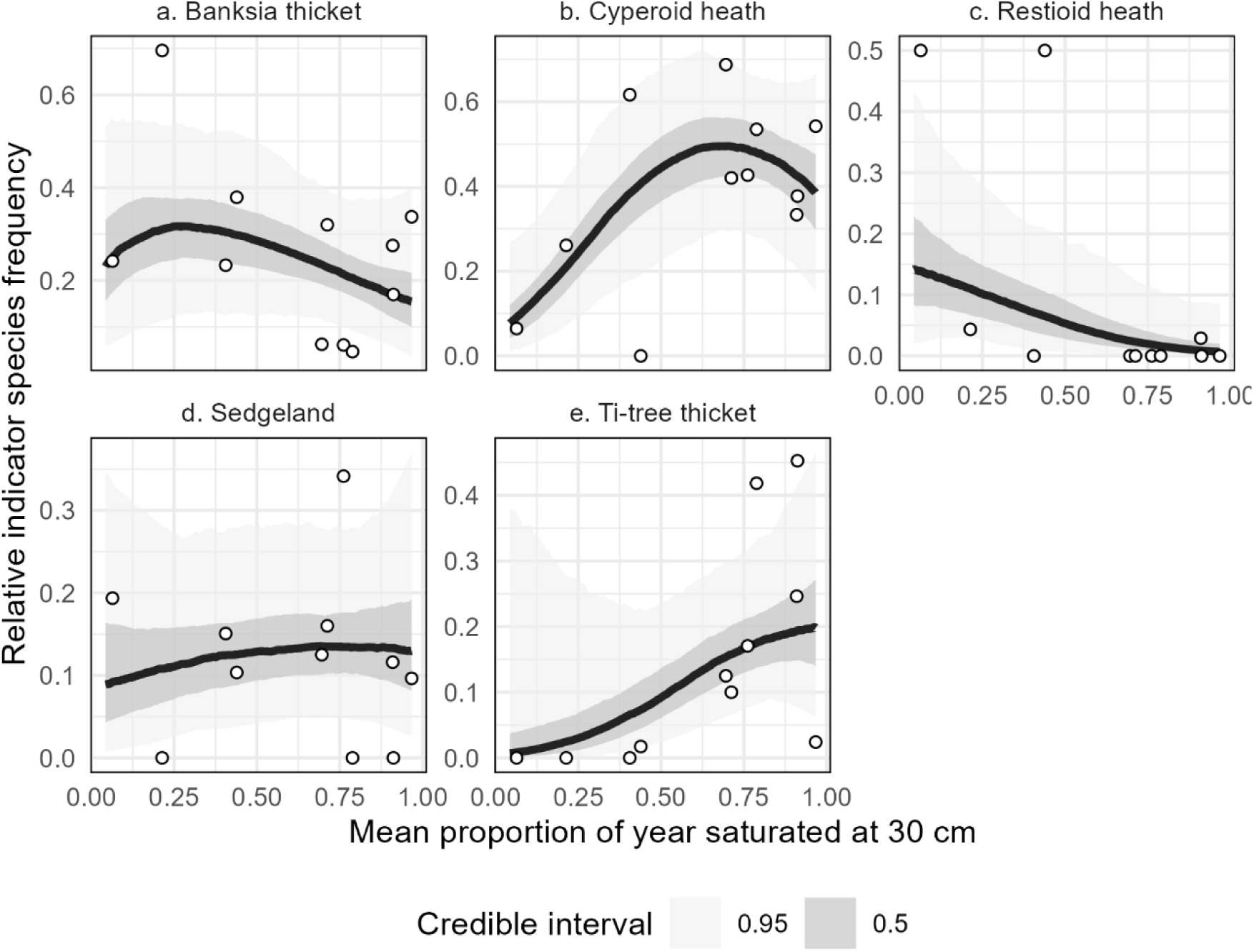
Modelled response of community indicator species relative frequency in five Upland Swamps to mean annual saturation at 30 cm depth. Points show observed frequency data while curves show the mean of the posterior predictive distribution with shading indicating 0.5 and 0.95 credible intervals.

Comparison of observed and predicted frequencies for indicator species in the three mined sites also showed some clear patterns (Table 2). Indicator species for the drier-adapted Banksia thicket was strongly overpredicted, while the wetter Cyperoid heath and Sedgelands were strongly underpredicted. Predictions for Restioid heath and Ti tree were more consistent with observed frequencies, although the latter had extremely low empirical values. The predictions suggest the community should be Banksia thicket, while the empirical data offer more support for a Sedgeland or Cyperoid heath community.

**Table 2.** Observed and predicted indicator species frequencies for the three mined sites. Observed IS frequency = empirical frequencies for indicator species for each community, Predicted IS frequency is the posterior predictive distribution for each community from the model shown in Fig. 2. Positive PPD = fraction of posterior predictive probability that exceeds zero when the observed mean is subtracted from the PPD. A reliable prediction would result in a value of 0.5, low values suggest the model underpredicted indicator species frequency (more dominance of indicators for this community than expected given the soil saturation data for the site), high values mean the predicted frequencies were higher than observed.

| Community | Observed IS<br>frequency<br>mean ($\pm 1$ SD) | Predicted IS<br>frequency<br>(median, 90% CI) | Positive PPD |
| --- | --- | --- | --- |
| **Banksia thicket** | 0.05 (0.00, 0.12) | 0.26 [0.12, 0.44] | **0.99** |
| **Cyperoid heath** | 0.31 (0.04, 0.57) | 0.10 [0.03, 0.21] | **0.02** |
| **Restioid heath** | 0.19 (0.02, 0.36) | 0.14 [0.05, 0.29] | 0.30 |
| **Sedgeland** | 0.44 (0.36, 0.52) | 0.09 [0.03, 0.23] | **0.01** |
| **Ti tree thicket** | 0.01 (0.00, 0.03) | 0.01 [0.00, 0.03] | 0.48 |

The simplified model estimating a collective hydrological niche for the three communities adapted to drier conditions (Fig. S3.1) was inferior to the five-community model, explaining only around half the variation in indicator species frequencies (Bayesian R^2^ = 0.25). The estimated hydrological niche for Ti-tree and Cyperoid heath indicator species were little affected, while the estimated optimum hydrological niche for pooled drier composite communities (Restioid heath, Banksia thicket and Sedgeland) was closest to the five-community estimate for Restioid heath at 20 [<16, 116] days.

## Discussion

### Use of indicator species in quantifying wetland community hydrological thresholds

An important step in the use of indicator species for evaluating the impacts of changing environmental conditions is ensuring the indicators respond to the factor they are hypothesized to represent (Lindenmayer and Likens 2011). Here, we found taxonomic community indicator species frequencies largely confirmed the established role of hydrology in structuring their respective plant communities (Keith and Myerscough 1993, Mason et al. 2017). In addition, we not only attain quantitative estimates for the duration of soil moisture saturation that differentiate the communities (i.e., their hydrological niche) but also test the validity of the indicators themselves in terms of their hydrological sensitivity.

Consistent with the established community indicator species framework for Upland Swamps, frequencies for Ti-tree thicket indicator species frequencies were higher under longer periods of root zone saturation than Cyperoid heath, while frequencies for Banksia thicket and Restioid heath implied adaptation to drier conditions (Keith and Myerscough 1993, Mason et al. 2017). The hydrological niche for Sedgeland indicator species was somewhat higher than expected for the community, but Sedgeland indicator species frequencies also showed the weakest response to saturation and the greatest scatter in the data (Fig. 2d). This suggests that while indicative of the community taxonomically, they are perhaps not as sensitive to the specific hydrological conditions under which the community is observed in nature (Keith and Myerscough 1993). One possibility is, that while fire differentiates Restioid heath, Sedgeland and Banksia thicket under drier conditions (Keith and Myerscough 1993, Mason et al. 2017, Mason et al. 2023), perhaps it is competitive hierarchies (e.g., for light) that allow Cyperoid heath and Ti tree thicket communities to dominate in wetter areas where sedgeland might otherwise develop. Alternatively, fire history at the sites might be a confounding influence on these results among the drier indicator species.

### Inferring thresholds of hydrology-driven change between communities

To plan suitable interventions to preserve wetland biodiversity, there is a need to understand how heterogeneous compositional forms emerge at both swamp and landscape scales (Janssen et al. 2024). The critical determinants of Upland Swamp vegetation communities are hydrology, driven by rainfall and groundwater saturation of the upper peat-forming layers (Cowley et al. 2019, Cairns et al. 2024) and the frequency of fire (Keith and Myerscough 1993, Mason et al. 2017, Mason et al. 2023). Focusing on hydrology, alignment with known patterns of occurrence in the field suggests the thresholds of saturation for the Cyperoid heath and Ti-tree thicket are probably reasonable estimates of their respective hydrological niche. Based on the estimated optima, a threshold of hydrological transition between Cyperoid heath and Ti-tree thicket communities would be expected to occur somewhere near 300 days average annual duration of saturation (i.e., near 0.8 in Fig. 2, Table 2).

Hydrological thresholds to the drier communities were less clear or consistent. For example, the estimated optimal saturation duration for Restioid heath communities was 16 - 29 days, which would be at the low end of durations for development of wetland sedge communities in Australia (typically at least 3 months duration; Roberts and Marston 2011). This might not prove a reliable estimate for the community overall. However, mid-point between the lower and upper estimates for Cyperoid heath and Banksia thicket (or the Drier composite community) respectively suggest somewhere near the 160-170-day mark (∼0.5) could be appropriate.

Some simple rules of thumb that emerge from the inferred community hydrological niche preferences above are that average annual root zone saturation of 1 - 6 months favours the three drier communities, 6-10 months most favours Cyperoid heath, while Ti tree thicket likely prefers periods of greater than 10 months. The least certain of these is the minimum threshold differentiating true terrestrial habitat and the drier communities, but this is a global change threshold that requires greater attention and specific approaches (Deane et al. 2025). Because of the limited number of sites available for this study, all thresholds should be viewed as preliminary, but they do offer a quantitative expectation for further hypothesis testing and contribute to understanding of the hydrological niche.

### Environmental water requirements and upland swamp ecosystems

The science of environmental water requirements originated in riverine systems, and is typically premised on the protection of key ecological flow bands within the natural flow regime (e.g., Poff et al. 2010). This approach is poorly suited to saturation driven systems such as Upland Swamps (Cairns et al. 2025). Similar ecosystems are found throughout Australia and internationally and, because of their hydrology, they are floristically diverse and support many ecotonal (or even terrestrial) species, often including threatened flora and fauna (Wilson and Paton 2004, Deane et al. 2016, Trezise et al. 2021). The estimated soil saturation durations, and thresholds between the communities offer a basis to couple projected change scenarios from Upland Swamp hydrological models to vegetation response (e.g., Cairns et al. 2024) and offer a further dimension to our understanding of their environmental water requirements (Cairns et al. 2025). Future work on environmental water requirements for Upland Swamps (and comparable vegetation communities) can use these estimates as null hypotheses for testing and refinement.

In cases where hydrological impacts occur in Upland Swamps, hydrological niche estimates herein can be used to predict the nature of the equilibrium vegetation community that develops post-impact. For example, in the three mined sites where indicator species frequencies were predicted in this study it was clear that 3-4 years post longwall mining extraction (see Mason et al. 2021) the frequency of Banksia thicket indicator species was lower than expected, while those for extant Cyperoid heath and Sedgeland indicators were clearly higher than expected. We predict that over time, one of the three drier communities will increasingly replace Cyperoid heath at these sites with fire expected to mediate equilibration of vegetation with the post-mining hydrological regime, the pace of change, and which community emerges. Periodic repeat vegetation monitoring at these sites could offer valuable insights into the duration of any time lag between the onset of hydrological impacts and the establishment of the new equilibrium plant community.

## Acknowledgements

We acknowledge all researchers who contributed to the establishment and monitoring of the long-term weather station network, particularly Christopher Simpson, Will Glamore, Adam McSorley and Duncan Rayner. This project was funded in part by the New South Wales Government through its Environmental Trust (Grants: RD 0134, RD 0028 and SSC 0049), DCD is funded by a grant from the Australian Research Council (DE240100477). Fieldwork was conducted under NSW Scientific Licence SL101124. We acknowledge the Dharawal people who are the traditional custodians of the land on which this study was conducted.

## Supporting information

Three appendices are provided as follows: Appendix 1 – de-trending and data cleaning for soil moisture data; Appendix 2 – Model diagnostics; Appendix 3 – Supplementary results

