## Supporting information for "Quantifying the hydrological niche of swamp vegetation communities using indicator species"

### Appendix 1 – de-trending and re-scaling of soil moisture data

Inspection of the soil moisture data offered clear evidence that the soil profile at 30 cm depth was saturating, indicated by either a more-or-less constant horizontal value over extended (Fig. S1.1a) or short-term periods, where it regularly reached the same maximum value (Fig. S1.2a). One of these two patterns was common to probes at all included sites but these presented two problems in using the data directly to determine the period of soil saturation at 30 cm depth.

First, was the drifting effect over time, where saturated observations fluctuated within and among years, typically following a linear increasing trend; second the probe reading at maximum saturation was universally less than 100%, despite clearly reaching a maxima (Fig. S1.1, S1.2).

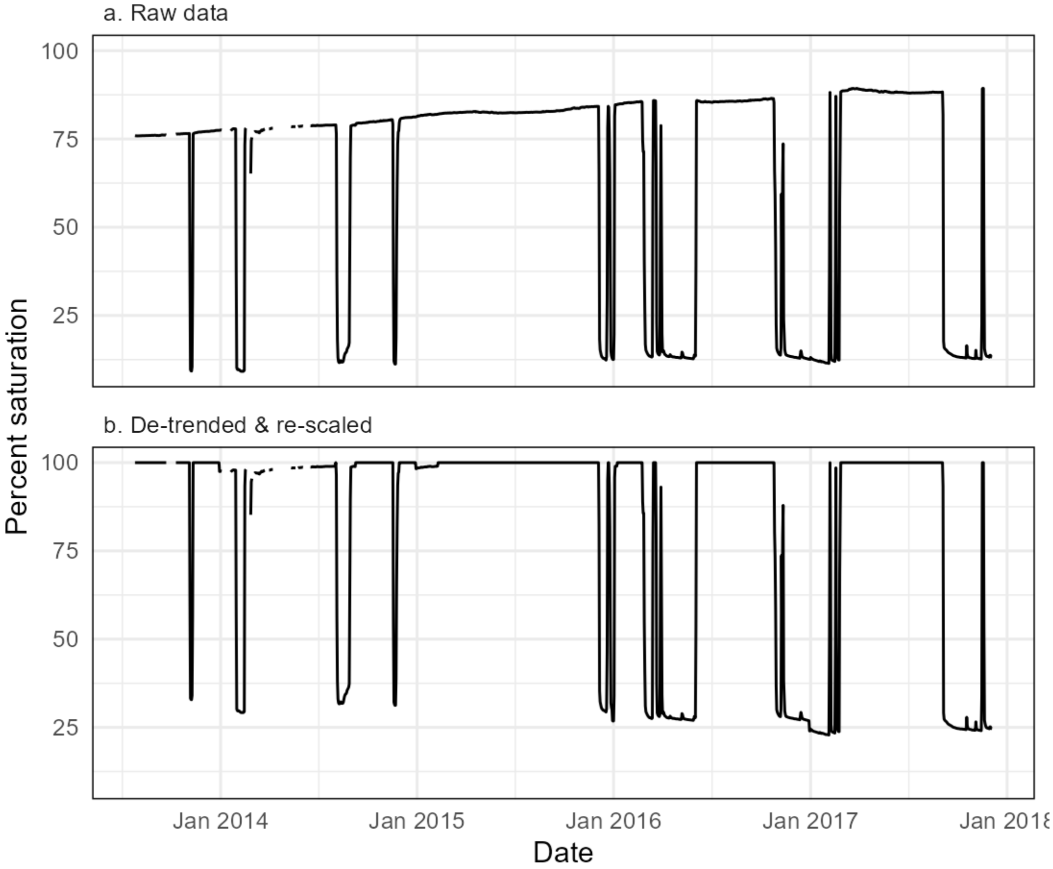


Figure S1.1. Example of the de-trending and re-scaling of soil moisture data for a site with evidence of long-duration saturation at 30 cm (Swamp A, probe 0). a. observed data from the soil probe, note clear evidence of saturation (horizontal records) and the change in maximum value over the period of record. b. de-trended data, after fitting quantile regression to the 99$^{th}$ data percentile, using the slope of the quantile regression line to remove the trend, then re-scaling the de-trended data to adjust the maximum observed value to 100% saturation.

To address these issues, we used a moving window approach to de-trend and re-scale data to a maximum of 100%. A non-overlapping local neighbourhood of 100 days was used in calculations, and the 99$^{th}$ percentile value of the raw data within the window was assumed to represent 100%. The difference between the 99$^{th}$ percentile and 100% was used to re-scale all values within the window to a maximum of 100%, with all values greater than 100% set to that value (i.e., to avoid saturation >100%). The corrected record was inspected to ensure the resulting curve reflected the dynamics of the raw data (see Fig. S1.1, S1.2).

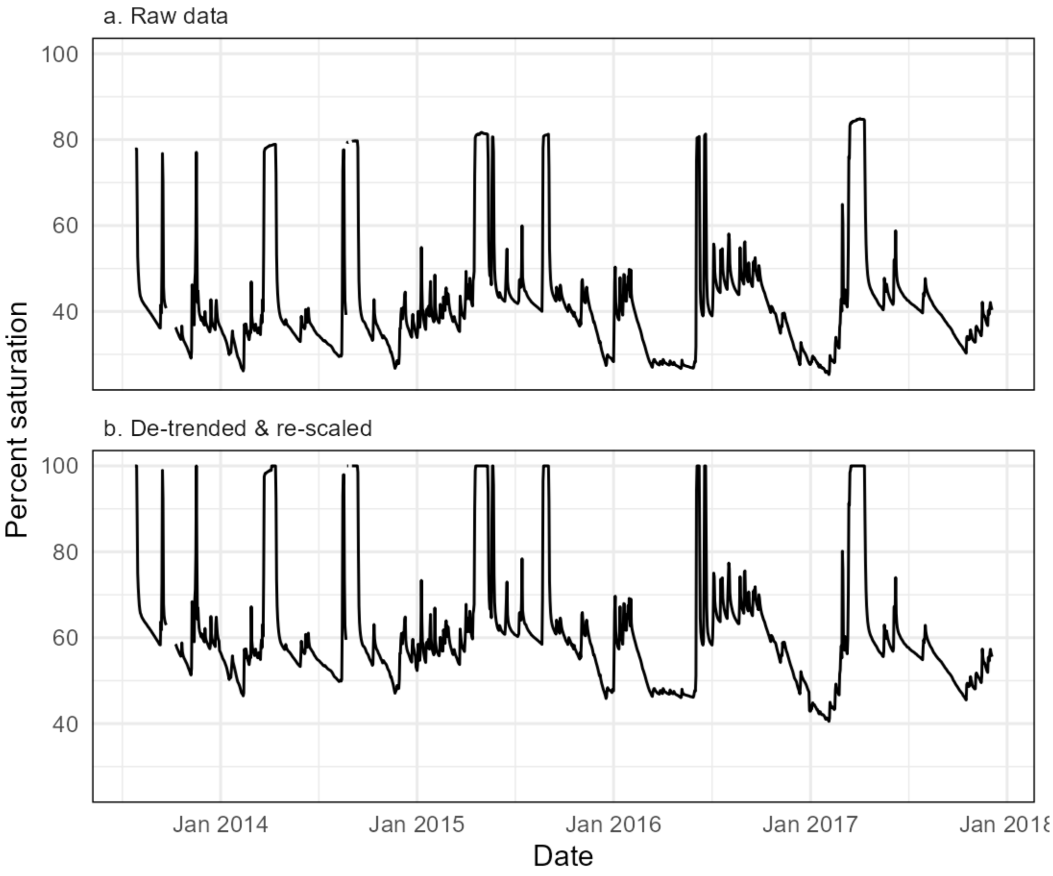


Figure S1.2. Example of the de-trending and re-scaling of soil moisture data for a site with only short duration saturation at 30 cm (Swamp 1b, probe 2). a. observed data from the soil probe, note clear evidence of saturation (horizontal records) and the change in maximum value over the period of record. b. de-trended data, after fitting quantile regression to the 80$^{th}$ data percentile, using the slope of the quantile regression line to remove the trend, then re-scaling the de-trended data to adjust the maximum observed value to 100% saturation.

**Appendix 2 - Model diagnostics**

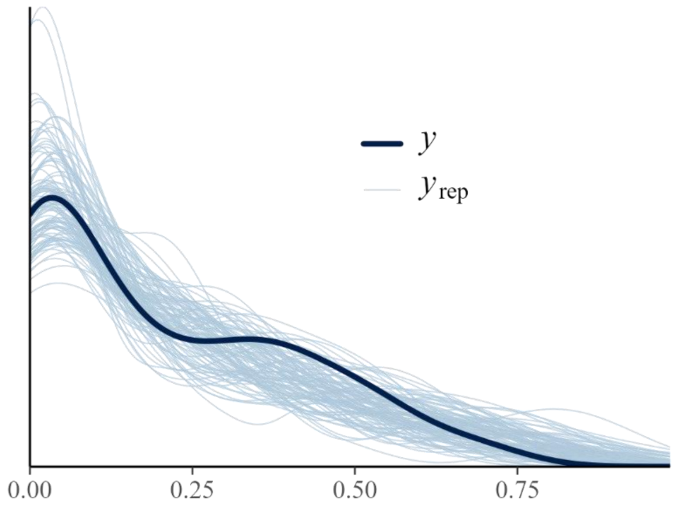


Fig. S2.1. Draws from the modelled posterior predictive distribution ($y_{rep}$) and empirical data distribution ($y$).

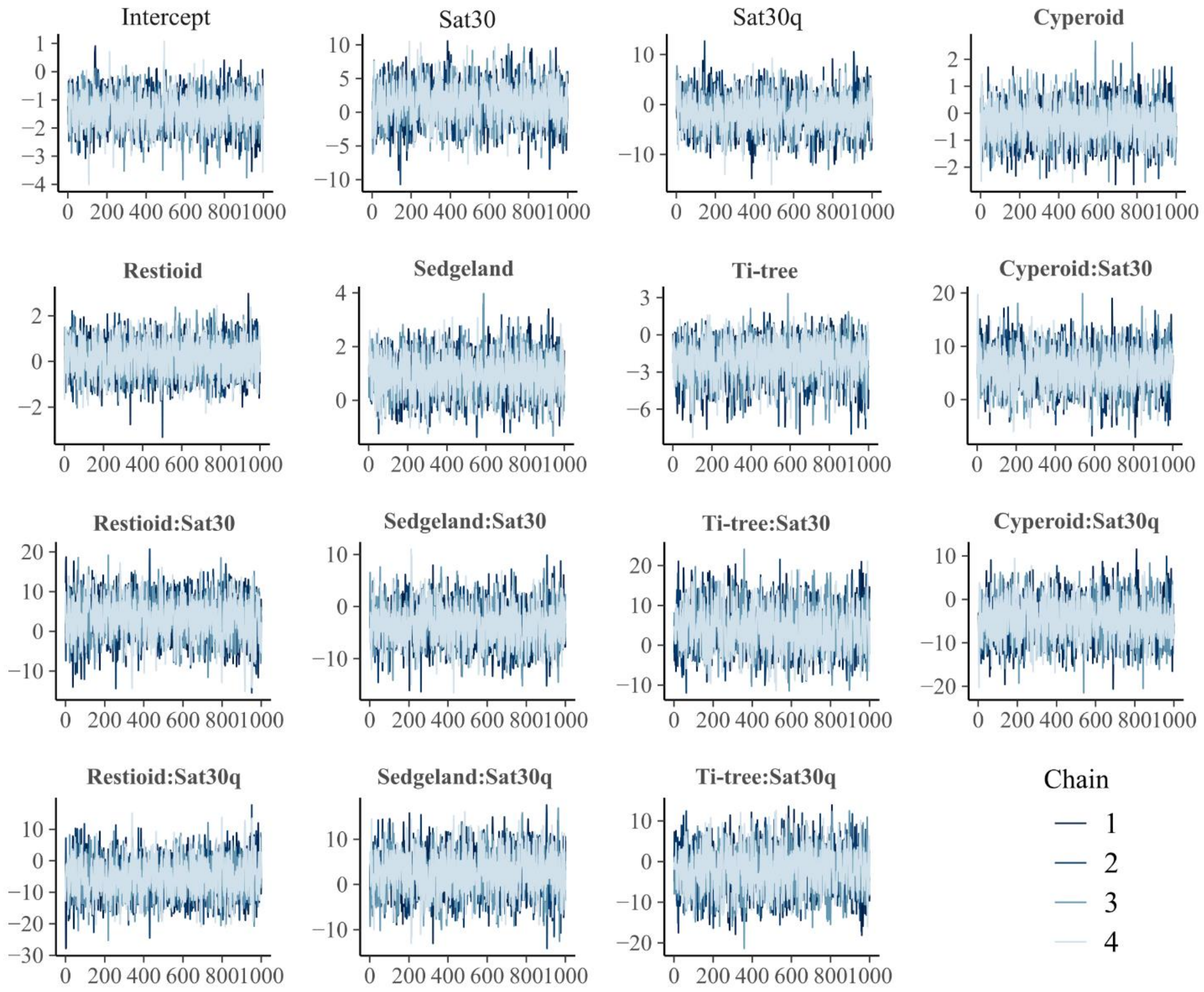


Fig. S2.2. Traceplots showing good mixing of the MCMC chains.

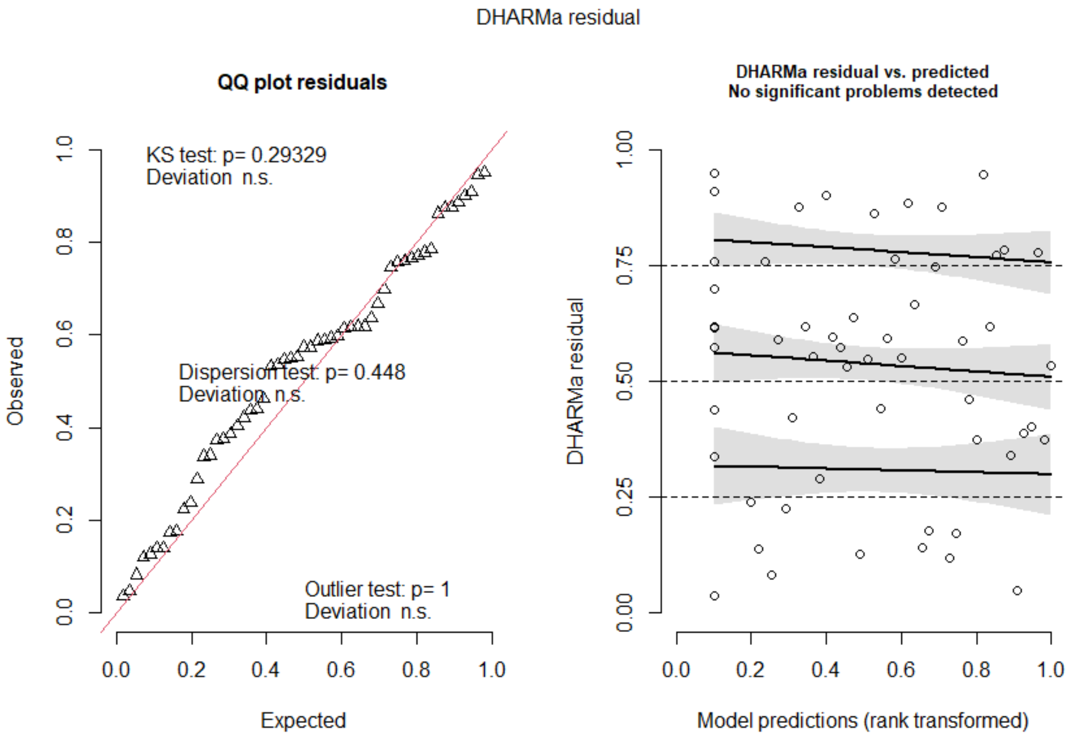


Fig. S2.3. Model residual tests from R package DHARMa (Hartig 2024) Left: QQ plot for uniformity of residuals; Right: residual vs fitted value quantile test.

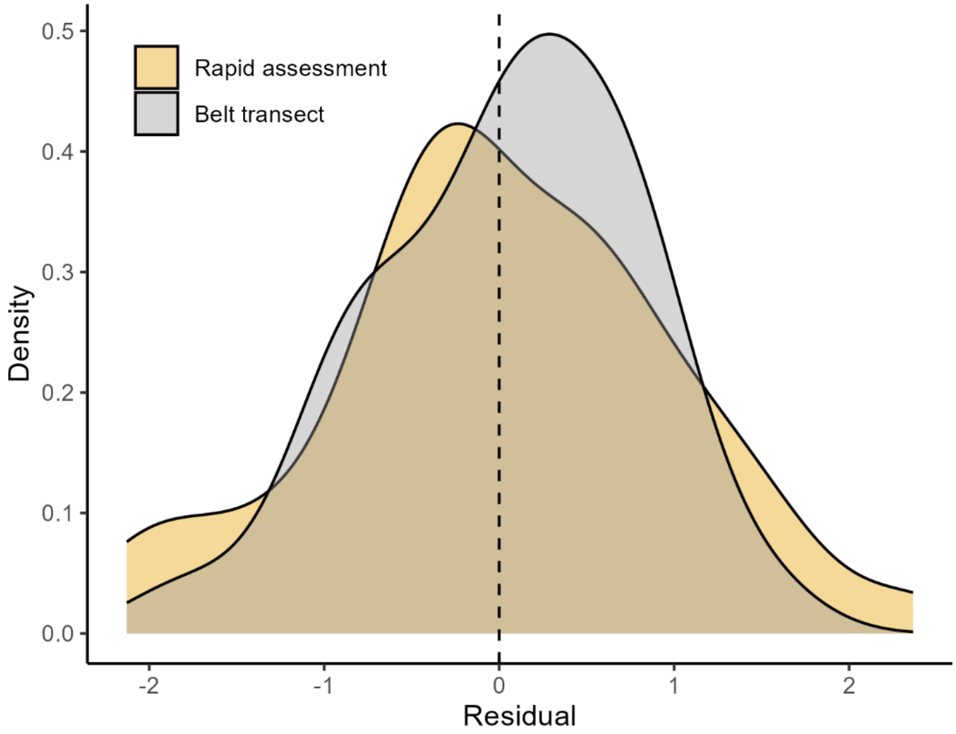


Fig. S2.4. Distribution of residuals according to survey method. There was no statistical evidence of a difference in median value (KW test, chi-sq = 0.05, *df* = 1, *p* = 0.83).

**Appendix 3 – Supplementary results**

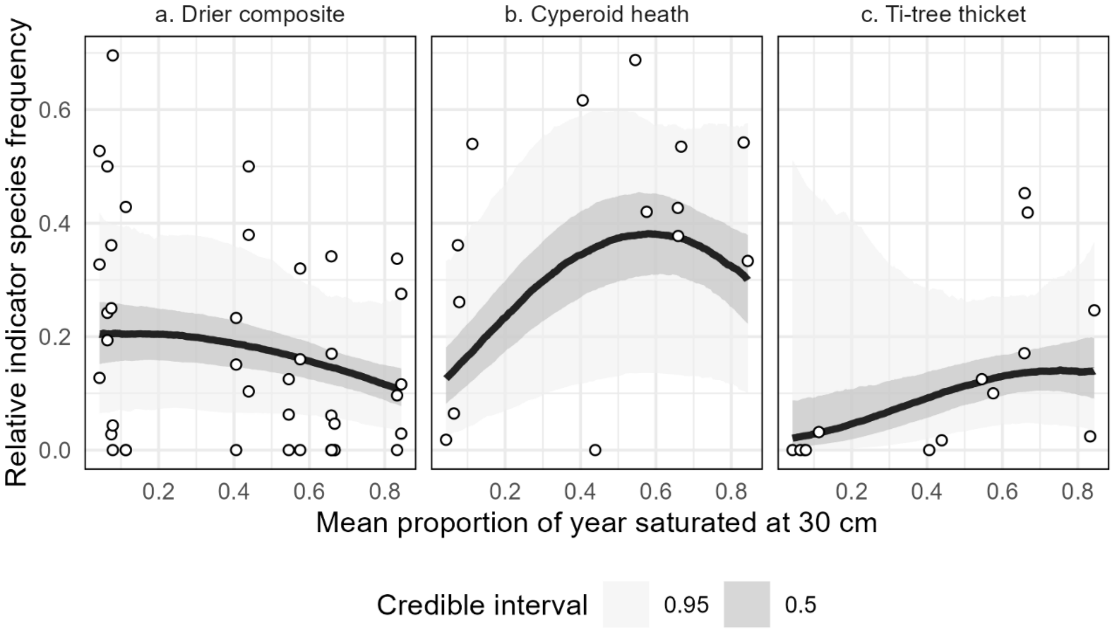


Figure S3.1. Modelled posterior predictions when the three dry-adapted communities are pooled. Here, "Drier composite" captures the pooled relative frequency of all indicator species for Banksia thicket, Restioid heath and Sedgeland. The hydrological response for the two communities adapted to wetter conditions (Panels b and c) were unchanged from the modelled results for all five communities. Compare Fig. 2, main text.
